# Bacteriophages from human skin infecting coagulase-negative *Staphylococcus:* diversity, novel species and host resistance

**DOI:** 10.1101/2023.11.07.565964

**Authors:** Samah E. Alsaadi, Hanshuo Lu, Minxing Zhang, Gregory F. Dykes, Heather E. Allison, Malcolm J. Horsburgh

## Abstract

The human skin microbiome comprises diverse populations that differ temporally between body sites and individuals. The virome is a less studied component of the skin microbiome and the study of bacteriophages is required to increase knowledge of the modulation and stability of bacterial communities. *Staphylococcus* species are among the most abundant colonisers of skin and are associated with both health and disease yet the bacteriophages infecting the most abundant species on skin are less well studied. Here, we report the isolation and genome sequencing of 40 bacteriophages from human skin swabs that infect coagulase-negative *Staphylococcus* (CoNS) species, which extends our knowledge of phage diversity. Six genetic clusters of phages were identified with two clusters representing novel phage species, one of which we characterise and name Alsa phage. We identified that Alsa phages have a greater ability to infect the species *S. hominis* that was otherwise infected less than other CoNS species by the isolated phages, indicating an undescribed barrier to phage infection that could be in part due to numerous restriction-modification systems. The extended diversity of *Staphylococcus* phages here enables further research to define their contribution to skin microbiome research and the mechanisms that limit phage infection.

## Introduction

Human skin is a complex and dynamic ecosystem that is colonised by diverse bacteria, fungi, and viruses that together form the skin microbiota (Grice *et al*., 2011). The composition and diversity of skin microbial communities is affected by multiple host and environmental factors, such as the topography and occlusion of sites, sex, age, and lifestyle (Grice *et al.,* 2009; Skowron *et al*., 2021). Despite variations in the microbial communities across different niches, the microbiota contributes functional roles collectively with resident skin cells towards maintaining skin homeostasis. In particular, the resident microbiota promote colonisation resistance that inhibits pathogenic organisms and regulates the immune system (Flowers *et al.,* 2020; Ochlich *et al*., 2023; Pastar *et al*., 2020; Severn *et al*., 2023).

Metagenomic studies have confirmed the abundance of viruses on human skin, which can impact its health (Hannigan *et al.,* 2015; Graham *et al*., 2022). This insight has increased focus upon the viruses of bacteria and the contribution to the skin virome of these bacteriophages (phages). Since phages infect and replicate inside bacterial hosts, they are proposed to have important roles in modulating the skin bacterial communities and their population dynamics (Hannigan *et al.,* 2015; Chevallereau *et al.,* 2022). Phages with a lytic life cycle can infect skin colonising bacteria, potentially modulating the composition of the bacterial community; this is exploitable for therapeutic applications (Liu *et al.,* 2015; Hatoum-Aslan, 2021). In contrast, temperate phages can integrate their viral genome into the bacterial host chromosome, and the resulting prophages can encode niche promoting activity, such as Sa3int phages of *Staphylococcus aureus* that are proposed to increase host colonisation, limiting their therapeutic usage (Rohmer and Wolz, 2021). Temperate phages comprise the major population of dsDNA phages sampled on skin and were found to be persistent (Hannigan *et al.,* 2015). Moreover, horizontal gene transfer by phage transduction promotes the evolution of a bacterial community through the acquisition of host accessory genes into genomes (Juhas *et al.,* 2009).

Coagulase-negative staphylococci (CoNS) naturally colonise human skin soon after birth with several CoNS species being present at differing abundances across the body. *Staphylococcus epidermidis* is the most frequently isolated CoNS and is considered a ubiquitous human skin resident (Otto, 2009). Other frequently isolated commensal CoNS of skin include *S. hominis*, *S. haemolyticus*, *S. lugdunensis,* and *S. capitis,* with the latter abundant on the scalp (Becker *et al.,* 2014; Hannigan *et al.,* 2015; Byrd *et al.,* 2018; Chong *et al.,* 2022). Although CoNS form part of the healthy skin microbiota, these staphylococci have the potential to cause opportunistic invasive disease, ranging from indwelling device infections to neonatal septicaemia (Natsis and Cohen, 2018; Heilbronner and Foster, 2020; Both *et al*., 2021).

Differences in the relative abundance of the commensal CoNS species within the skin microbiota could reflect their relative competitive fitness concomitant with ageing-related changes of the skin (Capone *et al*., 2011; Zhou *et al*., 2023). The range of bacterially produced factors that influence the abundances of staphylococci on skin are partially understood with studies showing the importance of antimicrobials and secreted antagonist proteins and peptides (Zipperer *et al*., 2016; Severn *et al.,* 2022a, 2022b). Beyond these antagonisms, improved understanding of microbiome dynamics and stability requires investigation of the effects of bacteria-phage interactions.

Consistent with the relative abundance of their hosts across different microenvironments, *Staphylococcus* phages are among the most abundant on skin (Oh *et al.,* 2016). Previous identification of skin phages used direct swabbing and purification of phages, and DNA extraction for metagenomic analysis (Hannigan *et al.,* 2015; van Zyl *et al.,* 2018). Most staphylococcal phages identified from skin have a linear dsDNA genome and belong to the class *Caudoviricetes* (previously order *Caudovirales* comprising families: *Siphoviridae*, *Podoviridae and Myoviridae*) (Hannigan *et al.,* 2015; Hatoum-Aslan, 2021). The majority of studies with staphylococcal phages have focused on *S. aureus,* particularly with respect to virulence genes and horizontal gene transfer (McCarthy *et al.,* 2012; Abatángelo *et al.,* 2017; Kizziah *et al.,* 2020). The study of CoNS and their phages has received less attention despite their interaction likely being a critical contribution to formation and temporal stability of skin communities and their microenvironment. Recent studies of staphylococcal phages include the isolation of lytic phage Andhra (Cater *et al.,* 2017; Hawkins, *et al.,* 2022) and multiple phages infecting *S. epidermidis* (Valente *et al.,* 2021), hinting at the potential for unexplored diversity on skin.

This study focused on isolating and characterising CoNS species-specific phages from the skin of healthy volunteers. A total of 40 phages infecting major CoNS species were purified from skin swabs. The phages were assigned to six genetic clusters revealing two clusters are previously undescribed phage species, of which we name one species as Alsa phage. We characterised the genetics and biochemical properties of four isolated Alsa phages. The collection of 40 phages were studied for their host range across 140 strains of 8 staphylococci revealing infection differences, establishing a basis for future studies on the role of staphylococcal phages in skin communities.

## Materials and methods

### Ethics approval

The study was approved by Central University Research Ethics Committee C at University of Liverpool, United Kingdom (project reference number: 9895).

### Bacterial culture

A collection of *Staphylococcus* species and strains collected by the Horsburgh lab group at the University of Liverpool was used in this study (Table S1). A single colony of each strain was inoculated in Brain Heart Infusion (BHI) broth (Neogen) and incubated overnight at 37 °C whilst shaking.

### Skin sample collection and treatment

Healthy volunteers (*n*= 80) were recruited to the study, and they used swabs to collect phages from their outer skin layer of different body sites. A total of 4 swabs were obtained from each participant. Swabbing was performed by soaking a sterile cotton swab (Copan) in sodium and magnesium salts (SM) buffer (100 mM NaCl, 4.8 mM MgSO_4_, 1 M Tris-HCl pH 7.5, 0.01% (w/v) gelatine) and rubbing the swab gently on the chosen skin site (∼5 cm x 5 cm) for a period of one minute. The skin swab samples were incubated overnight in BHI medium at 37 °C to enrich the phage yield of the collected samples. Cultured samples were then centrifuged at 5,000 rpm for 2 min to pellet bacterial cells and debris, and the supernatant was filtered through a syringe filter (0.45 μm, Star Lab) to obtain phage filtrate.

### Bacteriophage isolation

An agar-based spot assay was performed using the cultured skin sample supernatants to detect the presence of phages. Following the overnight incubation of *Staphylococcus* indicator host strains, 200 μl of mid-log phase cultured host cells were mixed with 4 ml of soft agar (0.4% w/v), then poured onto 1.5 % (w/v) BHI agar plates. Once the soft agar lawn solidified, a volume of 10 μl of phage filtrate was pipetted onto the bacterial lawn followed by overnight incubation of the plates at 37 °C. The presence of cleared growth zones in the indicator species lawn was used to pick plaques using sterile pipette tips to suspend in 500 μl SM buffer followed by filtering (0.45 μm).

### Bacteriophage purification and propagation

Plaque assays were performed to purify the isolated phages. Ten-fold serial dilutions of each isolated phage were prepared in BHI media supplemented with 10 mM MgSO_4_ (BHI-MG). Diluted phages (100 μl) were mixed with an equal volume of mid-log phase cells of susceptible bacterial strain and incubated at 37 °C with shaking for 1 h. The mixture was then added to 4 ml soft agar (0.4 % w/v), poured onto BHI agar plates (1.5 % w/v), and left to set prior to incubating overnight at 37 °C. A single plaque was picked into 500 μl SM buffer and filtered (0.45 μm) to obtain phage lysate. Plaque assays were repeated three times with filtrate to ensure purified phages.

To propagate phages, mid-log phase cells of susceptible bacterial hosts were infected with the purified phage lysates in BHI-MG and incubated at 30 °C for 18 h. After incubation, each phage propagation was centrifuged at 5000 *x g* for 5 min, the phage lysate supernatant was filtered (0.45 μm) and stored at 4 °C. Plaque assays were used to determine titres of the propagated phages.

### One-step growth curve

Bacteria for phage propagation were cultured in 10 ml BHI-MG to reach OD_600_ of 0.5 prior to addition of phage at a multiplicity of infection (MOI) of 0.1 followed by an incubation at 37 °C for 20 min to allow adsorption. Phage-adsorbed bacteria were collected by centrifugation at 8000 x *g* at 4 °C for 5 min. The supernatant was discarded, and cell pellets were re-suspended in 10 ml BHI broth and incubated at 37 °C for 90 min with agitation at 200 rpm. Aliquots of 200 μl were collected every 10 min and immediately serially diluted to measure phage titres using the soft agar overlay technique. The phage burst size was calculated by dividing the mean final titres by the mean initial phage titres.

### Lytic activity of phages

To assay host cell lysis by phages, an overnight culture of host bacteria was sub-cultured in BHI medium to OD_600_ of 0.2. Phages were added at a MOI of 0.01, 0.1, and 1. Bacteria with no phage added were used as a negative control. The mixture was incubated at 37 °C for 22 h, and OD_600_ was measured hourly. The OD_600_ measurements of all samples were plotted using GraphPad Prism (v9.3.1).

### Thermal and pH stability of phages

The stability of skin-isolated phages was assayed across a temperature range of −80 to +70 °C. One millilitre of phage solution (∼10^9^ PFU mL^-1^) was aliquoted in 1 ml of Phosphate Buffer Saline (PBS) and incubated at selected temperatures for 1 h, with control samples incubated at 4 °C. PBS adjusted to a range of pH 4 - 9 was similarly used to incubate phage solution at 28 °C for 1 h, with samples incubated at pH 7 as controls. Phage titres were determined using the soft agar overlay technique with ten-fold dilutions spotted on a lawn of its susceptible host bacteria. Statistical analysis was performed using analysis of variance (ANOVA).

### Host range determination

The host range of skin-isolated phages was assessed by testing each purified phage for its ability to infect 140 different strains belonging to 8 staphylococcal species (Table S1). This assay was conducted using the spot assay after overnight incubation, when the presence of clearing zones were visually categorised according to their clarity into three groups: complete, turbid, or zero.

### Efficiency of plating (EOP)

To validate the phage-host range data, an EOP assay was used with representative bacterial strains that showed complete or turbid clearing by spot assay (Table S1) against all infecting skin-isolated phages. To perform the assay, phage lysates were serially diluted to 10^-8^ dilution, and 10 μl of each dilution was spotted on the prepared bacterial lawns of each of the bacterial strains tested. The plates were then incubated overnight at 37 °C, the number of PFUs produced by the susceptible bacteria was counted, and phage titre was determined. EOP values were determined by dividing mean phage titres of a tested strains by the phage titre of the original propagation host.

### Phage adsorption assay

Exponential growth phase bacterial cells (∼10^8^ CFU mL^-1^) were infected with a volume of phage lysate to achieve MOI of 0.1 in a 10 ml BHI-MG. One millilitre of phage-host mixture was immediately collected upon phage inoculation to determine the initial phage titre. Phages were allowed to adsorb at 37 °C for 20 min and the same volume of the phage-host suspension was collected and centrifuged at 10000 x *g* for 3 min. Phages mixed with BHI only were used as a negative control. The titres of the free phages in the supernatants were determined by plaque assay. The proportion of unadsorbed phages was calculated by following the formula: [residual titre/ initial titre] X 100 %.

### Sodium periodate treatment

To determine the contribution of bacterial surface receptors in phage adsorption, treatment of host cells with sodium acetate (CH_3_COONa) (VWR International) and sodium periodate (NaIO_4_) (Sigma) was performed following previously published protocols (Sørensen *et al.,* 2011), with minor modifications. Following overnight culture of each strain, 500 μl of culture was centrifuged at 5000 x *g* for 5 min. Bacterial pellets were washed with PBS, re-suspended in 50 mM of NaIO_4_ in 50 mM of CH_3_COONa (pH 5.2), and incubated at room temperature in the dark for 1 h. Treated cells were centrifuged (10,000 x *g,* 5 min) and washed with 1 ml PBS. Pellets were re-suspended in 1 ml BHI broth and OD_600_ adjusted to 0.5. A phage adsorption assay with phages added at MOI 0.1 was performed as described above in the absence or presence of periodate treatment. Unpaired t-test was used for statistical analysis.

### Transmission electron microscopy

Washed phages from lysates were visualised by negative-staining using carbon film copper grids (Agar Scientific, UK; AGS160-3) charged to positive using a 2 min glow discharge in a Quorum Q150TS sputter coater with a turbo orifice plate. The phage samples were left to adhere to grids for 60 s in clamping forceps then grids washed with double distilled water and rinsed using 2% uranyl acetate, ensuring excess drops were wicked away with complete drying prior to imaging at 120 kV using a FEI120kV Tecnai G2 Spirit BioTWIN TEM with a Gatan Rio 16 camera plus digital micrograph software.

### Phage DNA extraction

Phage DNA was extracted following previously published protocols with some modifications (Jakočiūnė and Moodley, 2018; Rodwell *et al.,* 2021). To extract sufficient phage genomic DNA, phage particles were precipitated using precipitation solution of 10 % (w/v) polyethylene glycol (PEG) 8000 in 1 M NaCl. Samples were mixed gently by inversion and kept on ice for 2 h then incubated overnight at 4 °C. The precipitated lysates were centrifuged at 10,000 x *g* for 15 min. Supernatant was discarded and pellets re-suspended in 500 μl SM buffer. For phage DNA extraction, the phage mixture was treated with 20 U of DNase in DNase reaction buffer (Invitrogen), and 10 μl of RNase A (Invitrogen), mixed gently, and incubated at 37 °C for 1 h whilst shaking. This was followed by heat inactivation of the enzymes at 75 °C for 20 min. The purification of phage DNA was subsequently carried out using Norgen phage DNA isolation kit according to the manufacturer’s instructions. DNA concentration was measured fluorometrically using Qubit^TM^ dsDNA HS Assay Kit (Thermo Fisher Scientific). The integrity of each of the purified DNA was checked using 1 % Tris-Acetate EDTA (TAE) agarose gel electrophoresis.

## DNA sequencing and bioinformatic analysis

### Bacterial sequencing and MLST

DNA extraction of bacteria and Illumina sequencing analysis was performed by MicrobesNG (Birmingham, UK) using their established MicrobesNG Genome Sequencing Method (v20230314). The multi-locus sequence types (MLST) of strains were determined using MLST (v2.10) from genome sequences based on the MLST scheme of *S. hominis*. A core genome alignment was built from the *S. hominis* isolates and annotated using Prokka (v1.14.5) with default parameters. Pan-genome analysis of the isolates was done using Panaroo (v1.2.3) with default parameters producing the core genome alignment. The phylogeny of strains used for host range studies was constructed from a maximum-likelihood (ML) tree, with the substitution model TIM+F+I+R3 using 100 bootstrap replicates, generated by IQ-TREE (v.2.2.2.6) (Trifinopoulos *et al*., 2016) based on core genome alignment. The core genome maximum likelihood tree was visualised with iTOL (v6.8.1) (Letunic and Bork, 2021). The phylogeny of a larger set of *S. hominis* genomes (*n*=243) obtained from the NCBI database used the substitution model GTR+F+I+R10). Plasmidfinder (v2.0.1) (https://cge.food.dtu.dk/services/PlasmidFinder) (Camacho *et al.,* 2009; Carattoli *et al.,* 2014) was used to identify plasmids present in 26 *S. hominis,* 19 *S. capitis* and 25 *S. epidermidis* isolates with identity threshold of 85% and 80% coverage.

### Phage genome sequencing

MinION DNA library preparation of phage DNA was done prior to sequencing using Oxford Nanopore Technology (ONT) with the MinION sequencer. MinION sequencing libraries were prepared using Native Barcoding Expansion 1-12 (EXP-NBD 104) and 13-24 (EXP-NBD114) with the Ligation Sequencing Kit following the manufacturer’s instructions. Samples of the barcoded DNA library were mixed with sequencing buffer and loading beads then loaded onto a MinION SpotCN flow cell, version R9.4.1 using a MinION Mk1B device for sequencing using the MinKNOW (v21.06.0) software.

### Bioinformatic analysis of phage genomes

The generated raw dataset from the ONT was demultiplexed and base-called onto fastq files using Guppy (v4.5.4). NanoPlot (v1.32.1) and Seqstats (https://github.com/clwgg/seqstats) were used to check the quality of the sequencing reads. Reads were then *de novo* assembled using Flye (v2.9.2) pipelines. Quast (v5.0.2) was run to assess the assembly quality based on number of contigs, largest contig, total length, GC%, and N50. Assembly quality was also checked with Bandage (v0.9.0) and CheckV (v0.7.0). The genome ends (termini) and packaging mechanism were predicted and reordered using PhageTerm (v1.0.12) and tRNAs were searched using tRNAscan-SE (v2.0). PhageAI webserver platform (https://app.phage.ai/) was used to determine the predicted phage lifestyle. For each assembled phage genome, Pharokka (v1.0.0) was used for open reading frame (ORF) calling with annotation using default parameters combined with Prokka (v1.14.5) to ensure accurate determination of coding sequences (CDS). HHpred (https://toolkit.tuebingen.mpg.de/tools/hhpred) (Zimmermann *et al.,* 2018) was used to interrogate functional category determination by Prokka.

Family-level classification of phages was performed using ViPTree (https://www.genome.jp/viptree) using whole-genome derived amino acid sequences. In addition, the current inphared database (v1.2) was used to compare phage genomes using Mash (v2.3) and subsequently with whole-genome alignment using MUMmer (v4.0.0). For comparative analyses in this study, unverified phage genomes were excluded. To determine the relationships within each family, smaller phage genome datasets of each family were analysed by (i) calculating intergenomic similarities based on nucleic acid sequences using VIRIDIC (http://rhea.icbm.uni-oldenburg.de/VIRIDIC/) and (ii) constructing a phylogeny based on core proteins using intergenomic similarity thresholds for species (95%) and genus (70%). Core protein analysis of core genes was calculated with VirClust (http://rhea.icbm.uni-oldenburg.de/virclust/) and a tree cut at a distance of 0.9. Clinker (v0.0.28) (Gilchrist and Chooi, 2020) was used to perform the genome comparisons and describe relationships.

### Detection of anti-viral defence systems within bacterial genomes

Phage defence systems were detected with the PADLOC webserver (https://padloc.otago.ac.nz/padloc/) (v1.2.0) and padlocdb (v1.5.0) using genomes from strains of the host range analyses or *S. hominis* genomes retrieved from NCBI database. Defence systems categorised as DMS_other were excluded. In addition, the DefenseFinder webserver (https://defense-finder.mdmparis-lab.com/) (Tesson *et al.,* 2022) was used to detect anti-viral defence systems in CoNS strains. PHASTER (PHAge Search Tool Enhanced Release) webserver (https://phaster.ca/) was used to predict prophages of *S. hominis*, *S. capitis* and *S. epidermidis* strains as either intact, incomplete, or questionable.

### Statistics and reproducibility

For statistics and reproducibility in this study, each of the experiments were performed independently a minimum of three times.

## Data availability

All data supporting the findings of this study are available within the manuscript or supplementary information. The CoNS phage genome sequences are available as submission SUB13879236 (BioProject PRJNA1024390). The staphylococcal genomes are available as (BioProject PRJEB68213).

## Author contributions

SEA designed and performed experiments, characterised phages and performed DNA sequence and bioinformatic analyses. HL assisted in curating sequence data and contributed bioinformatic analyses. MZ assisted with phage characterisation. GFK performed EM. MJH and SEA conceived the study and with HEA designed experiments. SEA, HL, and MJH drafted the manuscript. SEA, HEA and MJH edited the manuscript.

## Results

### Isolation and identification of CoNS infecting phages from human skin

Skin was swabbed by healthy volunteers (*n*=80) to generate 320 skin samples that were processed and screened for the presence of phages infecting CoN *Staphylococcus* species. Isolation of phages involved the use of multiple strains of the staphylococcal species: *S. epidermidis*, *S. hominis*, *S. capitis*, and *S. haemolyticus,* with successive rounds of plaque purification. A total of 40 phages were isolated from the 320 skin swab samples and stored for study (Table S1). Purified phage genome DNA was sequenced using ONT to provide insights into phage diversity, and a phylogenetic tree was constructed based on the genome-wide sequence similarities compared against 1332 phage reference genome sequences (Figure 1). Comparative genome analysis assigned the collected phages to 6 genetic clusters, of which two are novel. Based on recent International Committee on Taxonomy of Viruses (ICTV) nomenclature (Turner *et al.,* 2023), each of the phages belong to an undefined order of the *Caudoviricetes* class. The novel phage genomes were assigned as unclassified members of the *Siphoviridae* family (cluster 4: genome length ∼45 kb) and an undefined *Herelleviridae* family (cluster 1: genome length ∼147 kb) (Figure 1). The previously identified phages included: undefined *Rountreeviridae* (genus *Andhravirus*) (cluster 2: genome length ∼18 kb); unclassified *Siphoviridae* (cluster 3 and cluster 5: genome lengths ∼46 kb and ∼40 kb, respectively); and unclassified *Sextaecvirus* (cluster 6: genome length 88-104 kb) (Table S1). The previously unreported *Herelleviridae* family (formerly part of *Myoviridae*) phage genomes are divergent from other *Staphylococcus* phage families identified from the skin sampling with maximal protein distance based on ViPTree analysis (Figure 1). These relationships between the isolated phages and their closest relatives were supported based on sequence homology, which determined variation across ascribed genera (Table S1).

**Figure 1:**
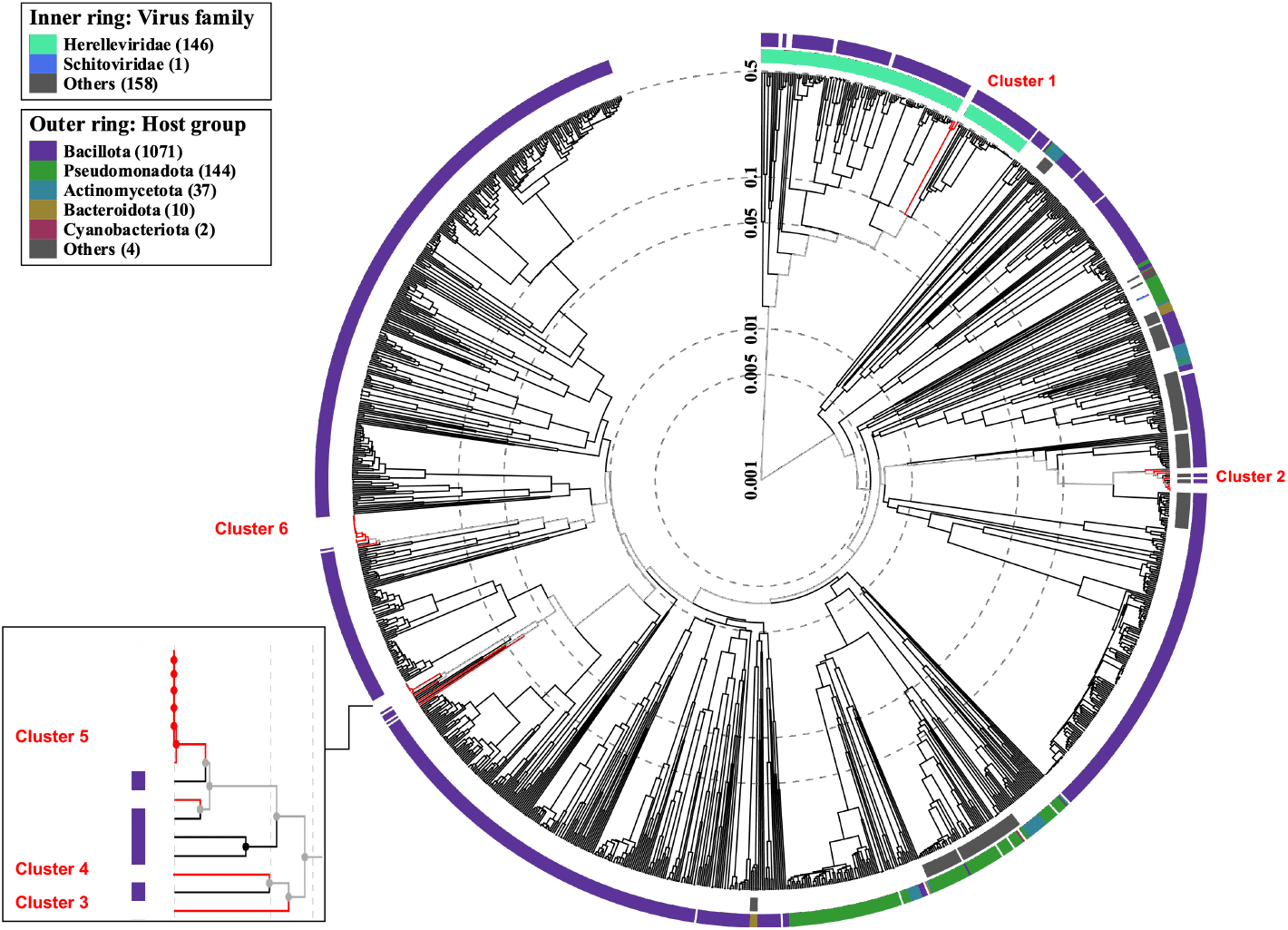
Phylogenetic tree of CoNS phages. Staphylococcal phage genomes from this study were assigned to 6 genetic clusters (1-6) using ViPTree 3.1 that included 1332 related phage taxa. Virus families are identified on the inner ring as *Herelleviridae* (mint green), *Schitoviridae* (blue), and other unknown families (grey). Phyla of their bacterial host is indicated in the outer ring with ascribed colours shown in the key. Cluster 4 represents a novel phage genome of this study (red line) and is distinct from the species øIME1354_01 representing the adjacent cluster of the same node (black line).

The staphylococcal phage genomes were compared using intergenomic similarities analysis (Figure S1) to further discriminate them based on their genetic properties by including the sequences of all 40 phages isolated in this study together with reference staphylococcal phage genomes (cluster 2: NC_047813 øAndhra; cluster 3: ON325435 øCUB-EPI_14; cluster 4: NC_070727 øIME1354_01; cluster 5: NC_028821 øStB2-like; cluster 6: NC_023582 øvB_SepS-SEP9). To validate the presence of distinct genetic clusters, phage genomes from ViPTree and inphared databases were included in the analysis based on the top k-mer match. Unverified phage genomes in GenBank were excluded from the analysis. The genome comparison supported the 6 identified clusters from the phylogenetic tree generated using ViPTree as being discrete, of which both clusters 1 and 4 represent new phage species genomes (Figure 1, Figure S1). The four cluster 1 *Herelleviridae* phage genomes are novel and named here as Alsa phages (type genome Alsa_1) based on their distinct clustering in the ViPTree output, which was supported from intergenomic similarities analysis (Figure S1) (Moraru, 2023).

Further discrimination of the 40 skin phage genomes and reference phage genomes was done based on intergenomic distances calculated by using VirClust hierarchical clustering of protein presence/absence. VirClust analysis highlighted the distinct protein content with 4 clades in the hierarchical tree showing limited overlap between the ascribed phage clusters (Figure 2). In addition to the Alsa phages discrimination, the singleton dsDNA phage (S-CoN_Ph26) genome of cluster 4 from intergenomic similarities analysis contained unique encoded protein clusters supporting it as a novel phage, and distinct from staphylococcal phage IME1354_01 (Tian *et al.,* 2022) representing the adjacent cluster of the same node as S-CoN_Ph26 (Figure 2, Table S1), but was not investigated further in this study.

**Figure 2:**
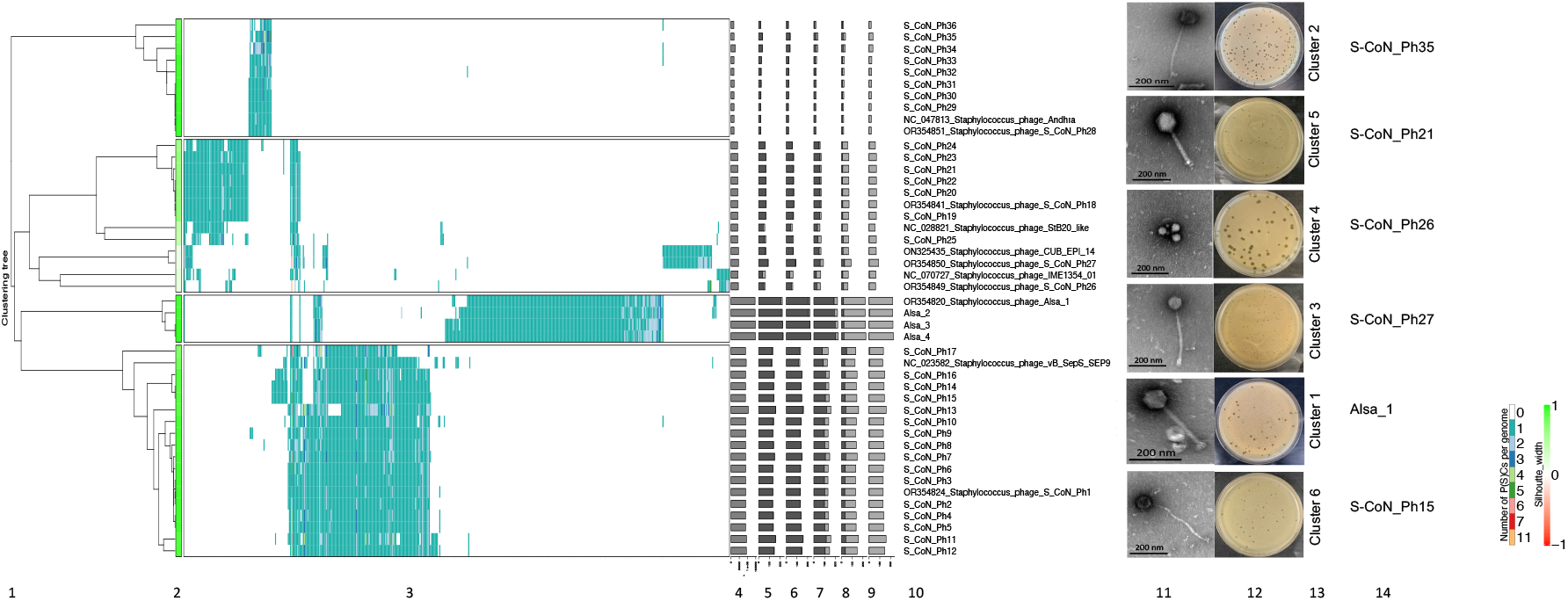
Hierarchical clustering of CoNS phage genomes. (1) Intergenomic distances of protein clusters were calculated using VirClust (2) and their relationship by silhouette width was examined using a hierarchical tree. The constituent viral genome clusters (VGCs) were determined based upon a 0.9 distance threshold. Similarity within each VGC is indicated by colour, with values closer to 1 being lime green. (3) Viral genome protein cluster (PCs) distribution is represented by a heatmap with rows representing individual genomes and columns as individual protein clusters together. A range of nine colours are used to indicate the number of protein clusters. Data to the right provide viral genome-specific statistics (4-9): (4) genome length (bp); (5) fraction of shared proteins (dark grey) from total proteins (light grey); (6) fraction of shared proteins within a VGC; (7) fraction of proteins specific to its own VGC; (8) proportion of proteins shared outwith a VGC; (9) the proportion of proteins shared exclusively outwith a VGC. (10) CoNS phage names from this study, provided as either their identifier or GenBank accession number, are included with reference phage genomes. (11) Electron micrographs of 6 phages, one from each cluster identified in this study [scale bar, 200 nm]. (12) Plaque morphology of each of the pictured phage and (13) their ViPTree cluster number and (14) their phage name.

The morphology of a representative phage of each of the 6 clusters defined by ViPTree analysis was determined by transmission electron microscopy (Figure 2). Phage S-CoN_Ph26 (cluster 4) has an icosahedral head (41 ± 0.4 nm) and a short, non-contractile tail (7.35 ± 0.4 nm) while the remaining phage cluster representatives are all long-tailed phages (Table S1). Analysis of the genome content using Phage AI predicted lytic lifecycles for 30 phages (clusters 1,2,6) and lysogenic lifecycle for the remaining 10 phages (clusters 3,4,5) (Table S1).

### Novel Alsa phages

Phage AI analysis was used to make predictions about the lifecycle of the Alsa phages identified in this study, which proposed they have a lytic lifecycle based on the absence of lambdoid phage integrase and repressor protein functions (Table 1, Table S1). Further phylogenetic analysis of staphylococcal phages identified that Alsa phages share a common ancestor with a broad group of lytic staphylococcal phages, including *S. aureus* øTwort and øK (Figure 3). The linear dsDNA Alsa phage genomes range in size from 146,115-148,384 bp, harbour 256-270 coding sequence (CDS), and feature a narrow G + C content ranging from 29.62-29.67% (Table 1, Table S1). The architecture of the four Alsa genomes shows synteny with collocated functional and structural proteins (Figure 4). No tRNA coding sequence was identifed using tRNAscan-SE and phageTerm identified a long Direct Terminal Repeat (DTR) sized from 8250-9667 bp (Table 1).

**Figure 3:**
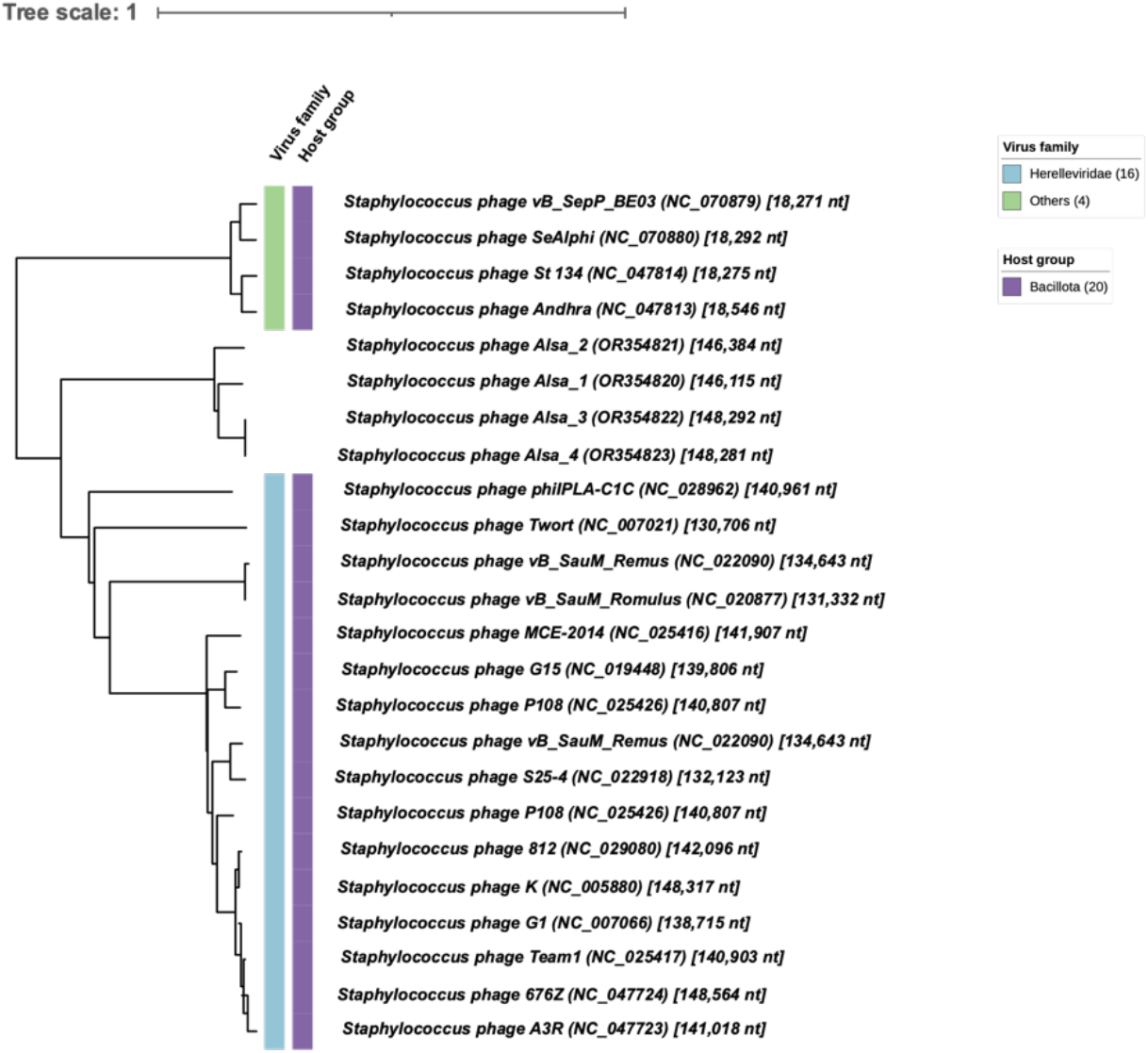
Phylogenic tree of Alsa and closely related *Staphylococcus* phages. Genomes of Alsa phages belong to the *Herelleviridae* virus family based on the tree generated by VIPTree 3.1 including closely related virus taxa calculated from genome distance matrix. Virus family taxonomy is indicated in blue (*Herelleviridae* infecting *S. aureus*) and green (other families) of the phylum Bacillota (host group).

**Figure 4:**
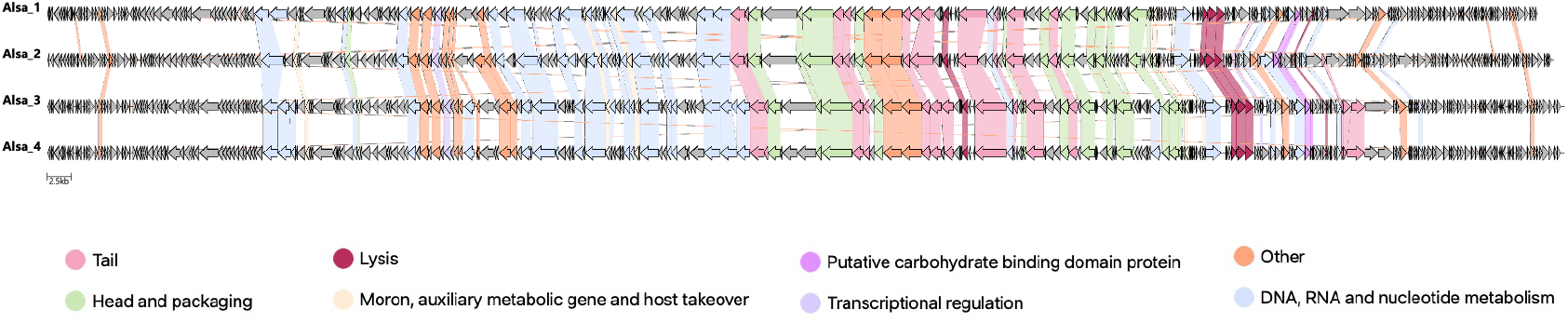
Comparison of Alsa phage coding regions based on functional annotation. Genomes of øAlsa1-4 were annotated using Pharokka and HHpred and visualised with Clinker with comparison of ORF functional categories across phages indicated by colour. Grey ORFs indicate hypothetical proteins. Scale, 2.5kb.

**Table 1:** General characteristics of Alsa phage genomes.

| Name | Alsa_1 | Alsa_2 | Alsa_3 | Alsa_4 |
| --- | --- | --- | --- | --- |
| Family | *Herelleviridae* |  |  |  |
| Predicted lifestyle | Virulent | Virulent | Virulent | Virulent |
| Total length (bp) | 148292 | 148281 | 146115 | 146384 |
| Number of contigs | 1 | 1 | 1 | 1 |
| GC % | 29.62 | 29.62 | 29.67 | 29.65 |
| CDS | 262 | 270 | 265 | 256 |
| Mean coverage | 891 | 826 | 892 | 715 |
| Direct Terminal Repeats (bp) | 8838 | 8839 | 8250 | 9667 |
| Class | DTR (long) |  |  |  |
| Packaging | COS |  |  |  |

### Characterisation of novel Alsa phages

One step growth curve assays were used to determine Alsa phage growth curves and their latent period. During replication, the latent period of both øAlsa_1 in *S. hominis* LIV1218 and øAlsa_2 in *S. hominis* LIV1220 was 15 min, while the latent periods of øAlsa_3 and øAlsa_4 in *S. capitis* 104 were longer at 20 and 30 min, respectively (Figure 5a). The burst sizes of the Alsa phages varied, ranging from 9 to 590 PFU per cell after 60 min, at least under the conditions studied. The doubling times of *S. capitis* 104, *S. hominis* LIV1218 and *S. hominis* LIV1220 were 1.17, 1.52 and 2.13 h, respectively. The thermal stability of the four phages across −80°C to 70°C was determined and revealed that øAlsa_1 and øAlsa_2 were stable at 50°C while both øAlsa_3 and øAlsa_4 had low stability at this temperature. Only øAlsa_1 and øAlsa_4 had stability at −20°C (Figure 5b). Each of the four phages were stable across a pH range from 4-9 (Figure 5c).

**Figure 5:**
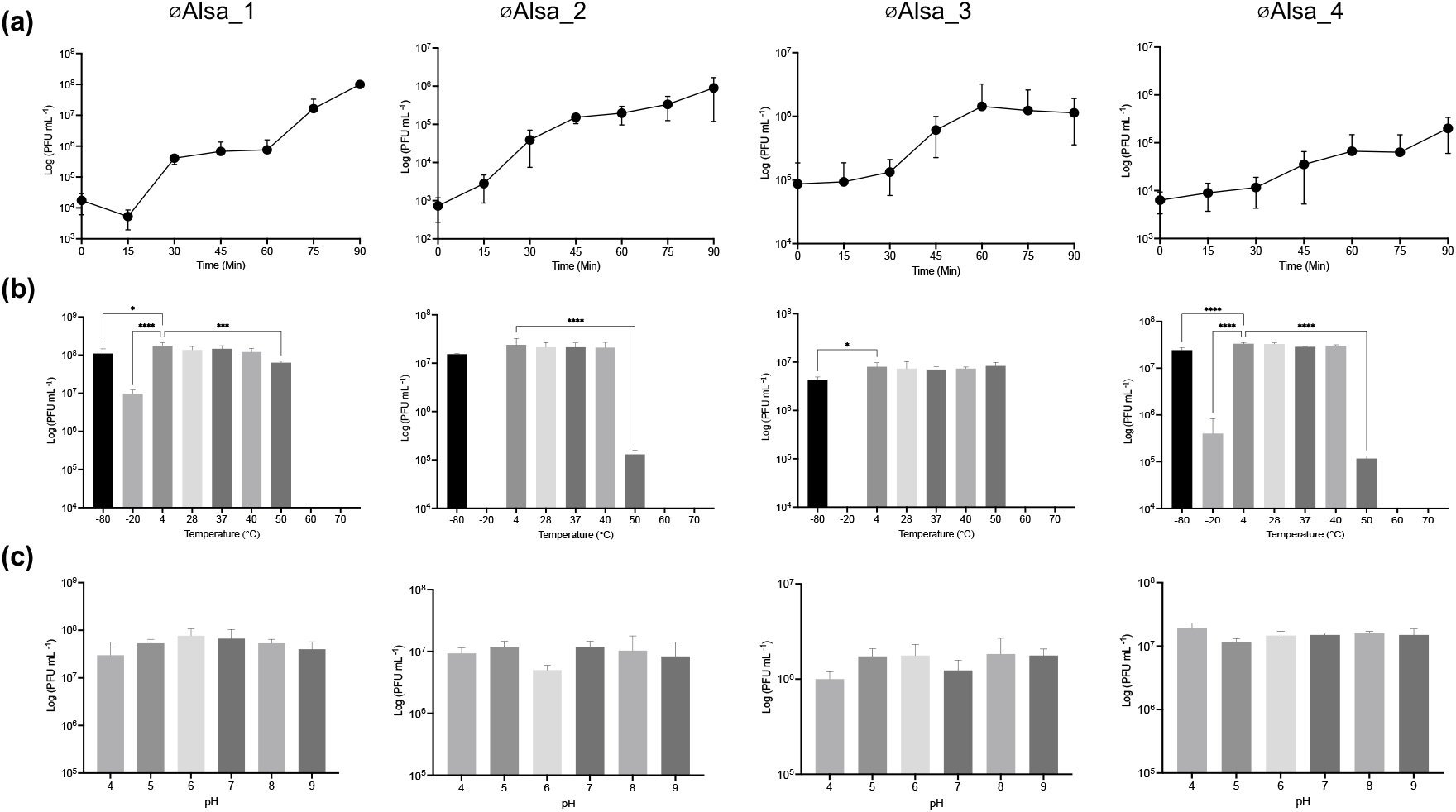
General characteristics of Alsa phages. One step growth curve of Alsa phages using original isolation hosts (a). Stability of phages across a range of temperatures (b) and pH (c) measured by PFU values. Statistically significant differences relative to control are indicated by asterisk: one-way ANOVA values * P<0.05, ** P<0.01, ***P<0.001.

The lytic activity of the Alsa phages was determined in their original host strains used for isolation with different MOI (0.01, 0.1 and 1) over 20 h in liquid culture. Relative to the absence of phage control, an MOI of 1 resulted in near complete lysis after 20 h. Distinct from the other Alsa phages, at MOI 0.1, øAlsa_4 did not cause complete lysis, while an MOI of 0.01 resulted in variable levels of lysis across the four phages in their respective hosts (Figure 6).

**Figure 6:**
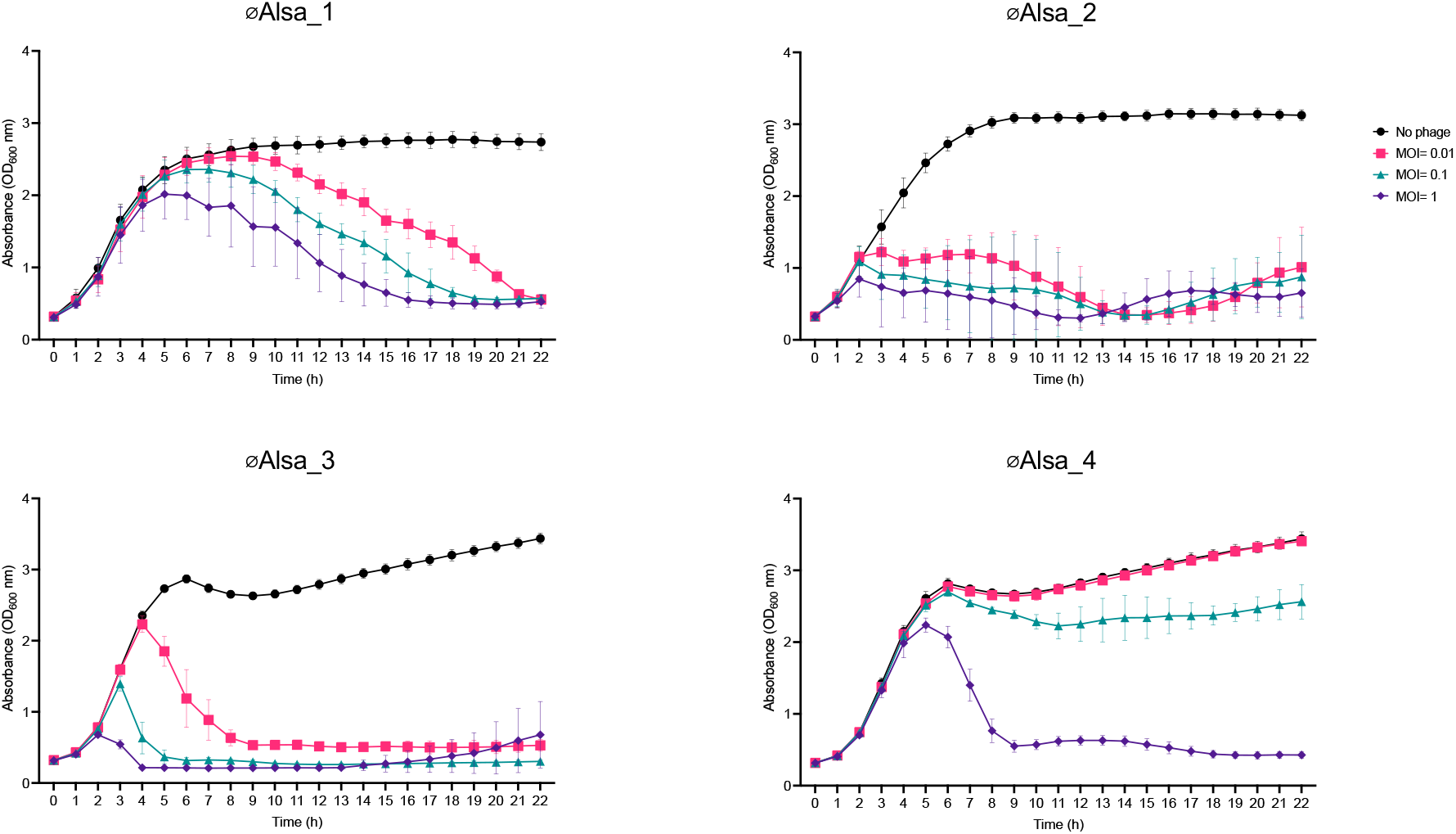
Lytic activity of Alsa phages. Phages were cultured with their respective isolation hosts at different MOI using the absence of phage as a control. Absorbance of cells was measured over 20 h in the absence (black, closed circle) or the presence of phages at different MOI (0.01, pink square; 0.1 green triangle; 1, purple diamond) with hosts LIV1218 (øAlsa_1), LIV1220 (øAlsa_2), 104 (øAlsa_3, øAlsa_4).

Adsorption of the four Alsa phages to their respective isolation strain hosts was determined using an MOI 0.1. Over a 20 min time period, between 70-100% of the population across the four phages adsorbed to the host cells, indicating high adsorption efficiency for each combination. To further characterise the binding of Alsa phages to their respective host, sodium periodate (NaIO_4_) treatment of the host cells, which would impact cell surface carbohydrate, resulted in significantly reduced adsorption efficiency of each phage compared to those incubated without NaIO_4_ (P= 0.001) (Figure 7). These data support Alsa phage binding to a host cell carbohydrate as a receptor with *S. capitis* and *S. hominis* differing in the extent of the reduction observed, which could reflect surface differences of these species.

**Figure 7:**
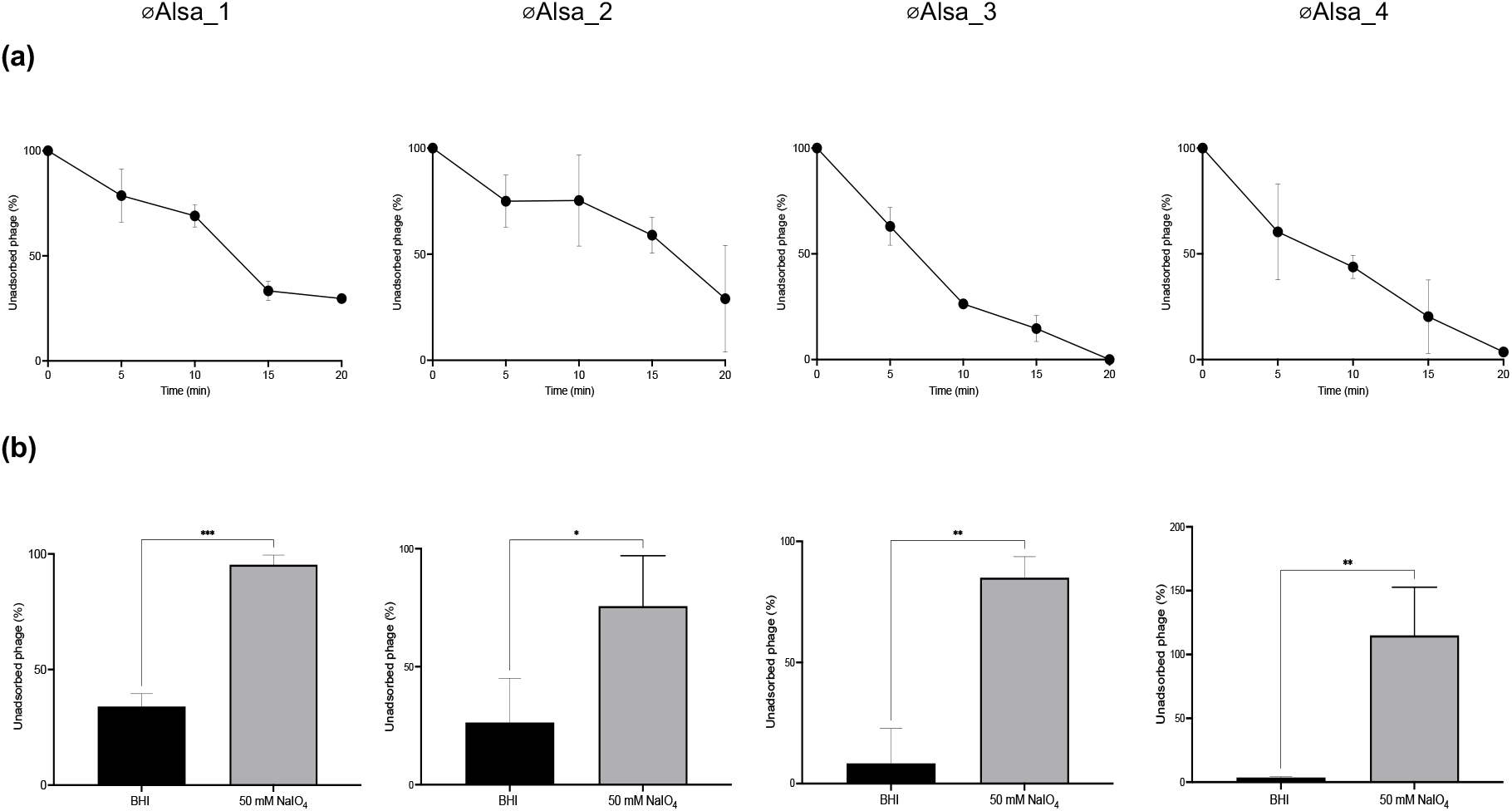
Adsorption of Alsa phages and effects of periodate treatment. (a) Alsa phages were incubated with their respective isolation hosts at MOI 0.1 and free phage were titred at the indicated time periods by plaque assay. (b) The effect of sodium periodate treatment for 60 min on the proportion of phages adsorbed (MOI 0.1) over 20 min was determined with hosts LIV1218 (øAlsa_1), LIV1220 (øAlsa_2), 104 (øAlsa_3, øAlsa_4).

### Host range determination of skin-isolated phages

To compare the host range of the 40 skin-isolated phages, a collection of 140 bacterial strains of 8 *Staphylococcus* species was used, including *S. lugdunensis*, *S. haemolyticus*, *S. epidermidis*, *S. capitis*, *S. hominis*, *S. warneri*, *S. saprophyticus*, and *S. aureus* (Table S1, Figure 8). Using a spot assay, the lytic activity of phages (titres 10^6^-10^9^ pfu ml ^-1^) was assessed after overnight incubation based on the extent of indicator strain lysis zones. *S. epidermidis*, *S. capitis*, and *S. warneri* were broadly susceptible to infection by the set of phages with individual strain variation (Figure 8). In contrast, *S. hominis* strains showed a distinct pattern of more limited infection by the set of phages, when compared to other *Staphylococcus* species tested (*S. epidermidis, capitis, warneri* and *lugdunensis*) (Figure 8, Figure S2). Within the set, the Alsa phages showed a capability for complete lysis of *S. hominis*. Specifically, øAlsa_1, øAlsa_2, øAlsa_3 and øAlsa_4 infected 9%, 8%, 5% and 3% of the 140 *Staphylococcus* strains tested while 50%, 42%, 15% and 27% of *S. hominis* strains (*n*=26) were infected, respectively.

**Figure 8:**
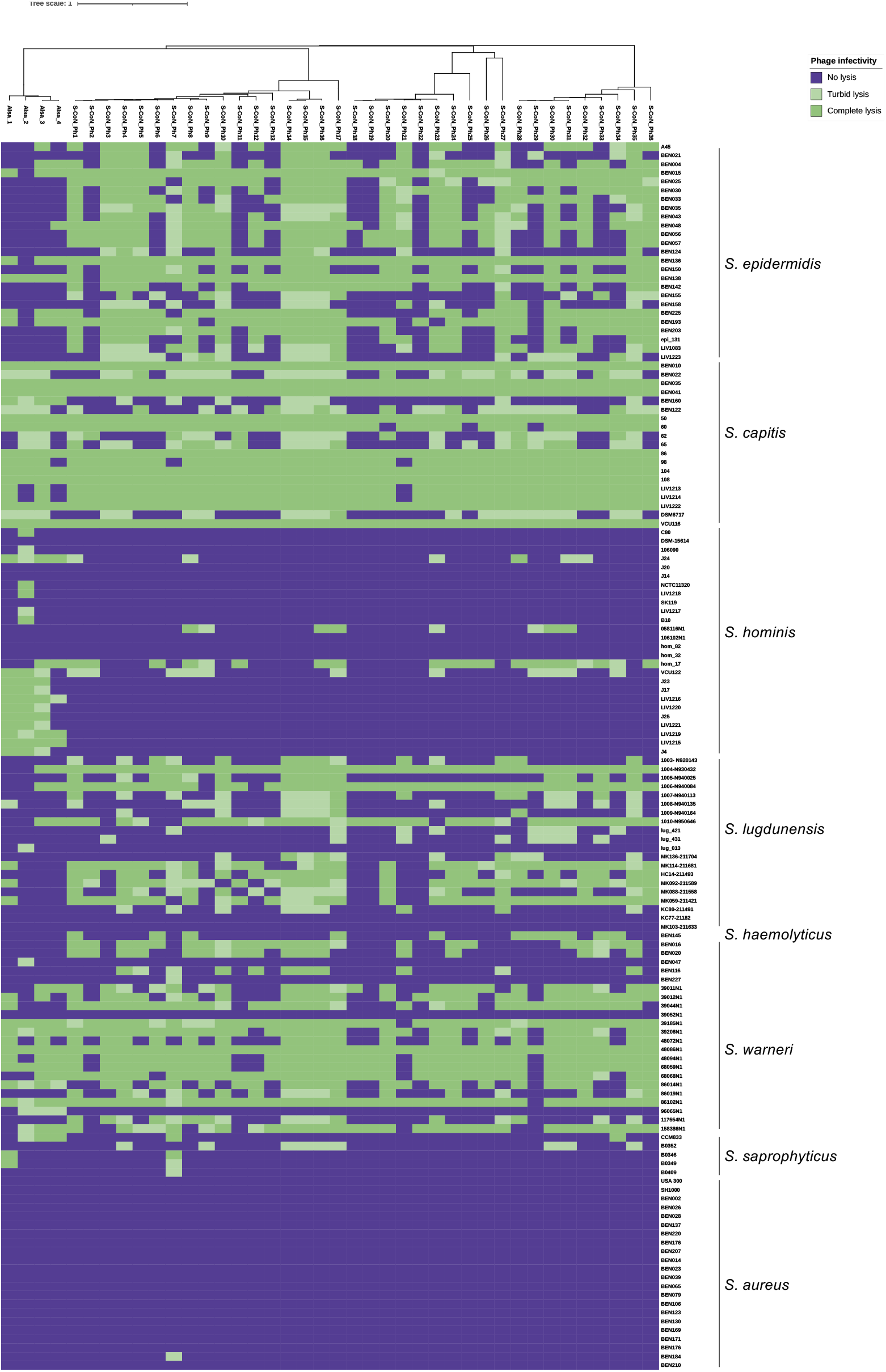
Host range of isolated CoNS phages. The 40 CoNS phages of this study were tested for their ability to infect 140 strains of staphylococcal species using a spot assay. Host lysis was determined visually by zone clarity with complete (dark green), turbid (light green), or no lysis (purple). The experiment was repeated a minimum of three times.

Further analysis of the included genome sequences revealed diversity of the *S. hominis* strains with 15 different sequence types (ST) identified with ST94 and ST96 (Figure 8, Table S1) having the greatest sensitivity to phages. Several of the phages in the identified set were more capable of lysis of individual CoNS species, specifically S-CoN_Ph30 caused complete lysis of 80% of the tested *S. epidermidis* strains (*n*= 20) while the phages S-CoN_Ph5, 11, 17, 20, 24, 28, 34 caused complete lysis of 79% of the tested *S. capitis* strains (*n*= 15). Distinct from the CoNS species, *S. aureus* was uniformly resistant to infection by all the tested phages. A single phage (S-CoN_Ph7) was capable of lysis from without, however no infection occurred as confirmed by the absence of plaques on *S. aureus* BEN184 upon serial dilution of the phage titre, and the inability to propagate anything from the infection supernatant on the original host strain, *S. capitis* 104 (Figure 8).

EOP assays were used to confirm phage amplification and plaque formation and validate the host restriction patterns observed with spot assays. Phages that caused complete or turbid lysis were selected to perform EOP assays with 7 representative host strains of both *S. epidermidis* and *S. capitis* species, together with 13 *S. hominis* strains (Figure 9). Broadly, the EOP data confirmed the distinctly reduced infection of *S. hominis* strains that contrasted with the greater infection of both *S. capitis* and *S. epidermidis* observed in the host range data, based on plaque formation (Figure 9, Table S1).

**Figure 9:**
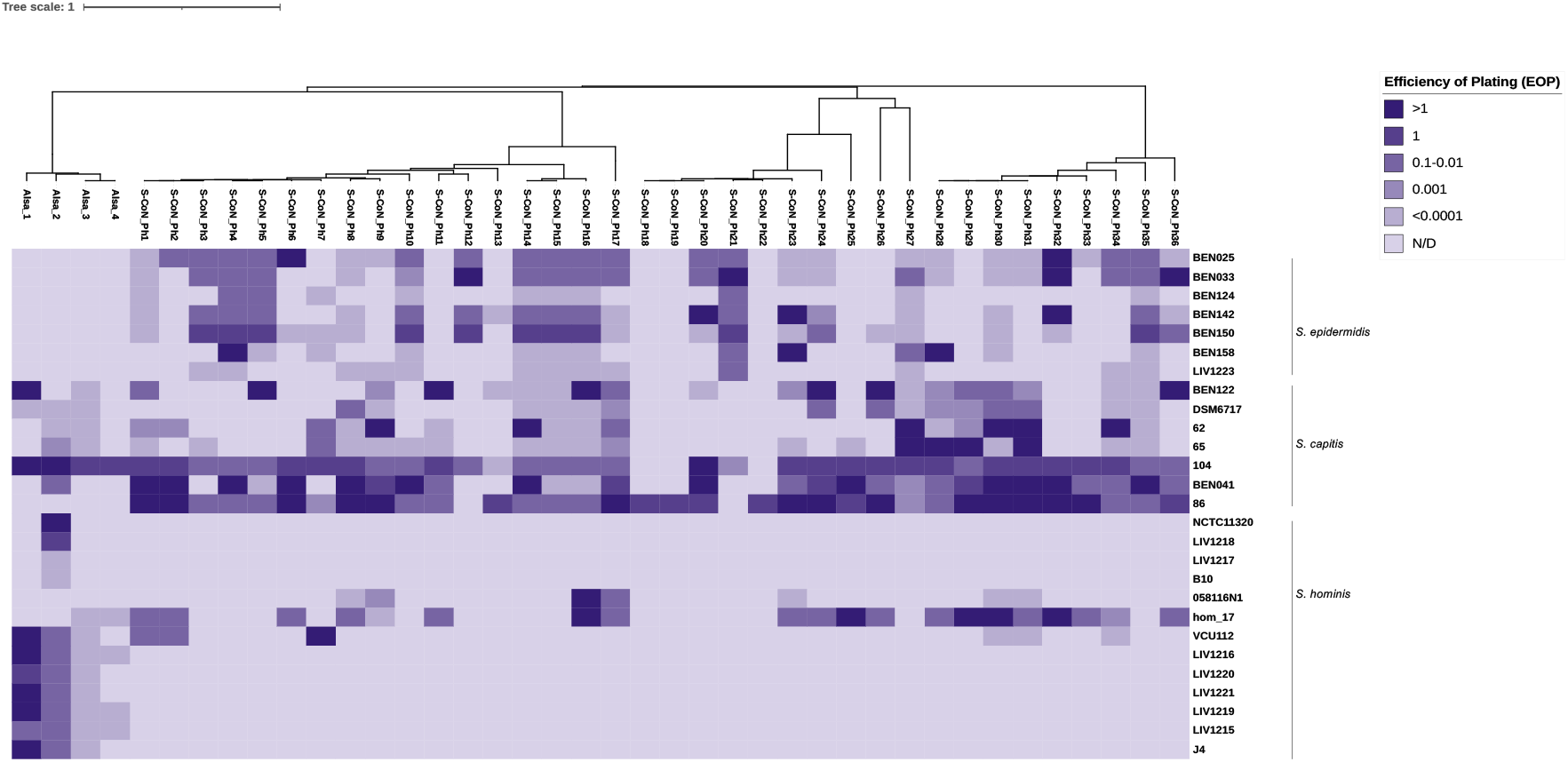
EOP assays of phages using hosts *S. epidermidis*, *S. capitis* and *S. hominis.* Representative strains of each host that showed complete lysis in the spot assay were tested for their ability to produce plaques, as calculated by PFU. EOP values were calculated relative to PFU values from propagation in their original hosts. This assay was repeated independently three times.

### Phage defence identification

To gain insights of potential defence mechanisms that could explain the limited phage infection observed here with *S. hominis*, the genomes of all the strains tested (*n*=26) were interrogated using both PADLOC (Figure 10) and DefenseFinder (Figure S3). Each of these defence mechanism identification analyses revealed no single mechanism of *S. hominis* to explain the host range phenomenon, although multiple restriction-modification (RM) systems were variably present. Subsequently, 243 genomes of *S. hominis* available in the NCBI database were analysed to look for defence systems more broadly at species level (Figure S4), again revealing the presence of multiple RM systems as a potential explanation for the limited infection. Comparative genome analysis of the *S. hominis* isolates tested with phages here identified that the number of intact prophages ranged from 0-1 with additional incomplete prophage regions (Figure 10) showing evidence of temperate phage infections. This indicates that among possible explanations, the phage infection barrier observed here could be a superinfection mechanism determined by a resident prophage, but this was not studied further. Plasmid content was also determined in each host strain and no clear link was identified between number of plasmids present and phage infection susceptibility (Figure 10, Table S1). Defence mechanism identification was also performed with the *S. capitis* (*n*=19) and *S. epidermidis* (*n*=25) genomes of strains included in the host range experiments (Figure S5, Figure S6). Different patterns of defence mechanisms were revealed to be present in each species genomes yet neither the presence of RM nor abortive infection mechanisms identified offered an unambiguous explanation for the host restriction observed in *S. hominis*.

**Figure 10:**
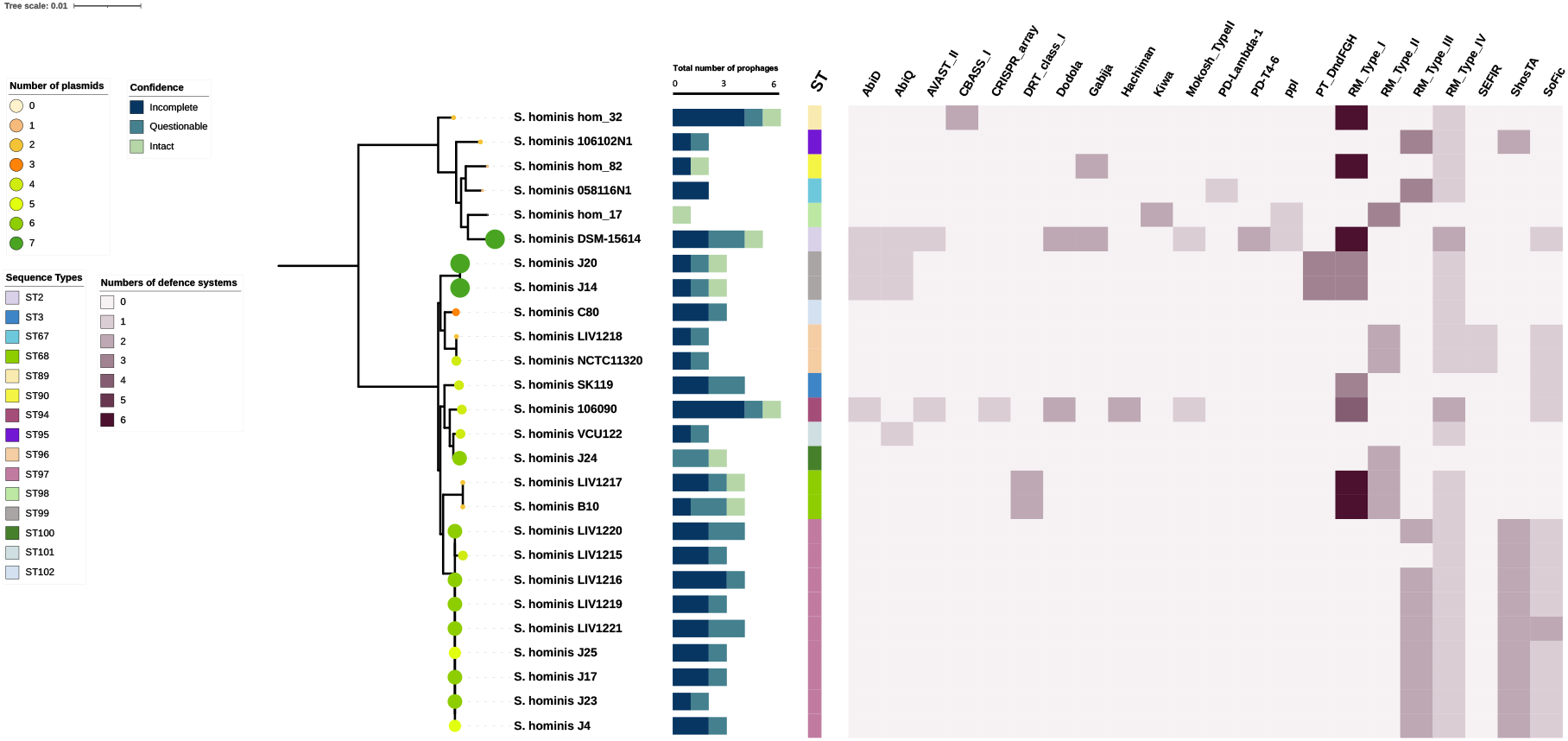
Phage defence systems and plasmid content of *S. hominis.* The presence of phage defence systems in genomes of the *S. hominis* strains used in the host range study were detected with PADLOC. Number of defence systems present is scored (0-6) and the total number of prophages (0-6) are indicated as either intact, incomplete, or questionable. Phylogenetic tree of *S. hominis* strains was generated using IQ-TREE. Sequence type (ST) of strains is indicated by vertical-coloured bars. Plasmid content (0-7 plasmids) is shown by different sized and coloured circle symbols, determined by PlasmidFinder.

## Discussion

Recent analyses have demonstrated the spatio-temporal distribution of bacterial species on human skin over the life course (Grice *et al.,* 2009; Byrd *et al.,* 2018; Flowers *et al.,* 2020). While other studies sought to determine the host and bacterial factors that drive microbial community changes, here we investigated the virome with respect to phages that infect CoNS. The contributions of both phages and their hosts is needed to improve understanding of the biotic and abiotic drivers of population changes on skin and the ability to externally alter these for health and personal care.

For this study, we focused on isolating phages present on human skin that infect the multiple coagulase-negative species of *Staphylococcus* that are abundant residents, where most efforts have focused previously on the skin pathogen *S. aureus*. We identified 40 phages from skin swabs of 80 volunteers that were isolated from different body sites.

Comparative genome analysis revealed that the phages isolated in this study formed six distinct genetic clusters that belong to an undefined order of the *Caudoviricetes* class (Turner *et al.,* 2023). Within this undefined order, two of the genetic clusters were previously reported and here we identified an additional two novel clusters, which extends our understanding of skin phage diversity with respect to CoNS species. Specifically, the novel phages belong to an unclassified *Siphoviridae* family (cluster 4: genome length ∼45 kb) and an undefined *Herelleviridae* family (cluster 1: genome length ∼147 kb). The latter of these phages named Alsa represents a clade that shares a common ancestor with known lytic *Herelleviridae* phages of staphylococci, based upon a phylogenetic tree produced by ViPTree (Figure 3). We propose that Alsa are lytic phages based on their relationship with reported *Staphylococcus* lytic phages together with the absence of integration-related genes. Further study is needed to investigate the proposed novel temperate phage S-CoN_Ph26 of cluster 4 that was not investigated further here. The successful isolation of this large collection of phages using simple and common swab sampling methods highlights the potential to uncover yet more diversity of the cutaneous virome via phages that infect staphylococci. Such studies should combine greater numbers of volunteers from distinct global populations with additional CoNS species for phage propagation.

The broadening of phage diversity from skin builds on several other studies that examined phages capable of infecting CoNS hosts. A recent study of 26 volunteers found half harboured phages that infect *S. epidermidis* and revealed 7 unique phage sequences (Valente *et al.,* 2021). An extensive screen of wastewater samples in Switzerland (Göller *et al*., 2021) uncovered 94 novel phages infecting 29 *Staphylococcus* species comprising 117 strains. Notably, the wastewater screen identified many phages infecting species *S. vitulinus, S. sciuri, S. xylosus, S. succinus, S. epidermidis* and *S. aureus*. Further analysis of the skin phages sampled in our study is needed to determine if their host range extends to those species in the wastewater study that are uncommonly found on skin. Hannigan *et al.,* (2015) reported that from cutaneous sample metagenomic data a relative proportion of >85% temperate phages, similar to that identified in the human gut virome (Reyes, *et al*., 2010; Minot *et al*., 2011). In contrast, our sampling size of 40 phage genomes identified a proportion of 25% temperate phage, which might reflect differences in the sampling procedure and chosen hosts for propagation.

The host range here with the 40 phages determined that the species *S. epidermidis, S. capitis*, *S. lugdunensis* and *S. warneri* were broadly infected, with a limited number of individual strains of these staphylococci showing an absence of infection that likely reflects their phage resistance (Figure S2). Within the collection of phages isolated here, we determined the host range was restricted to CoNS species with no evidence of infection leading to propagation in *S. aureus*. Further study is required to understand the barriers to these phages infecting *S. aureus* that might reflect the absence of phage receptors and could extend to phage defence mechanisms. Most phages of *Staphylococcus* adsorb to wall teichoic acid glycoploymers and recent work by Beck *et al*., (2023) identified that resistance to øE72 infection in *S. epidermidis* is mediated by WTA modification with glucose. The authors propose that glucose-modified WTA alters susceptibility to phage infection in certain CoNS species.

A clear pattern of reduced infection was evident with *S. hominis* using the isolated collection of phages, whereby few individual bacterial strains were infected (Figure S2). This reduced infection pattern of *S. hominis* compared with other tested CoNS species indicates there might be a widespread mechanism to limit phage infection in *S. hominis*. Contrastingly, Alsa phages were capable of directing the complete lysis of a greater proportion of *S. hominis* strains than other tested phages. Consequently, Alsa phages show a promising capacity to infect *S. hominis* and should be investigated further to exploit this property. Given that *S. hominis* is associated with human axilla body odour (Bawdon *et al*., 2015; Troccaz *et al.,* 2015), future studies should investigate whether there is a correlation between reduced presence of Alsa phages in high body odour individuals. Potentially, increasing infection by resident Alsa phages could be promoted to limit odour. For example, an inhibitor of the phage infection barrier could be used in personal care that would promote the resident skin phages to infect and reduce the skin population of *S. hominis*. Studies are required to determine the phage receptor of *S. hominis* and its likely glycosylation modifications. The periodate treatment used here supports that an extracellular carbohydrate could serve as the phage receptor for this bacterial species.

We investigated the potential mechanism of *S. hominis* phage resistance by curating possible phage defence mechanisms in our strain set (*n*=26) using PADLOC. In addition, 243 genomes available in public databases were analysed. Both strain sets revealed that multiple RM systems are widely present in *S. hominis*, including Type IV RM that was frequent in both the limited strain set tested in this study and in a wider curated publicly available strains set. Comparing with the *S. epidermidis* and *S. capitis* strains that showed widespread phage infection in this study, multiple RM systems were present in the genomes of both species, indicating no clear link between RM systems and reduced phage infection observed. Further investigation is needed to determine the exact molecular basis of the limited *S. hominis* infection by phages based on infection, adsorption and receptor studies, abortive infection systems and potential prophage mediated superinfection exclusion. Such interrogation will help to unravel aspects of the virome and the factors that could structure skin communities, and their contribution to health and disease.

The extended collection of phages infecting *Staphylococcus* in this study adds to the potential to enhance the molecular toolkit needed to improve recombinant DNA studies of the CoNS, which has lagged behind that of *S. aureus*. In addition, studies investigating use of the phages in reducing staphylococcal biofilms and in therapeutic and personal care applications can be addressed in future work.

## Supplementary

**Figure S1-S6**

**Table S1**

## Acknowledgements

We thank Tobi Somerville for providing staphylococcal isolates and Paul Loughnane for technical support. SEA was funded by a PhD scholarship from Taif University, Kingdom of Saudi Arabia.

