## Supplementary Figures S1-S6 for "Bacteriophages from human skin infecting coagulase-negative *Staphylococcus:* diversity, novel species and host resistance"

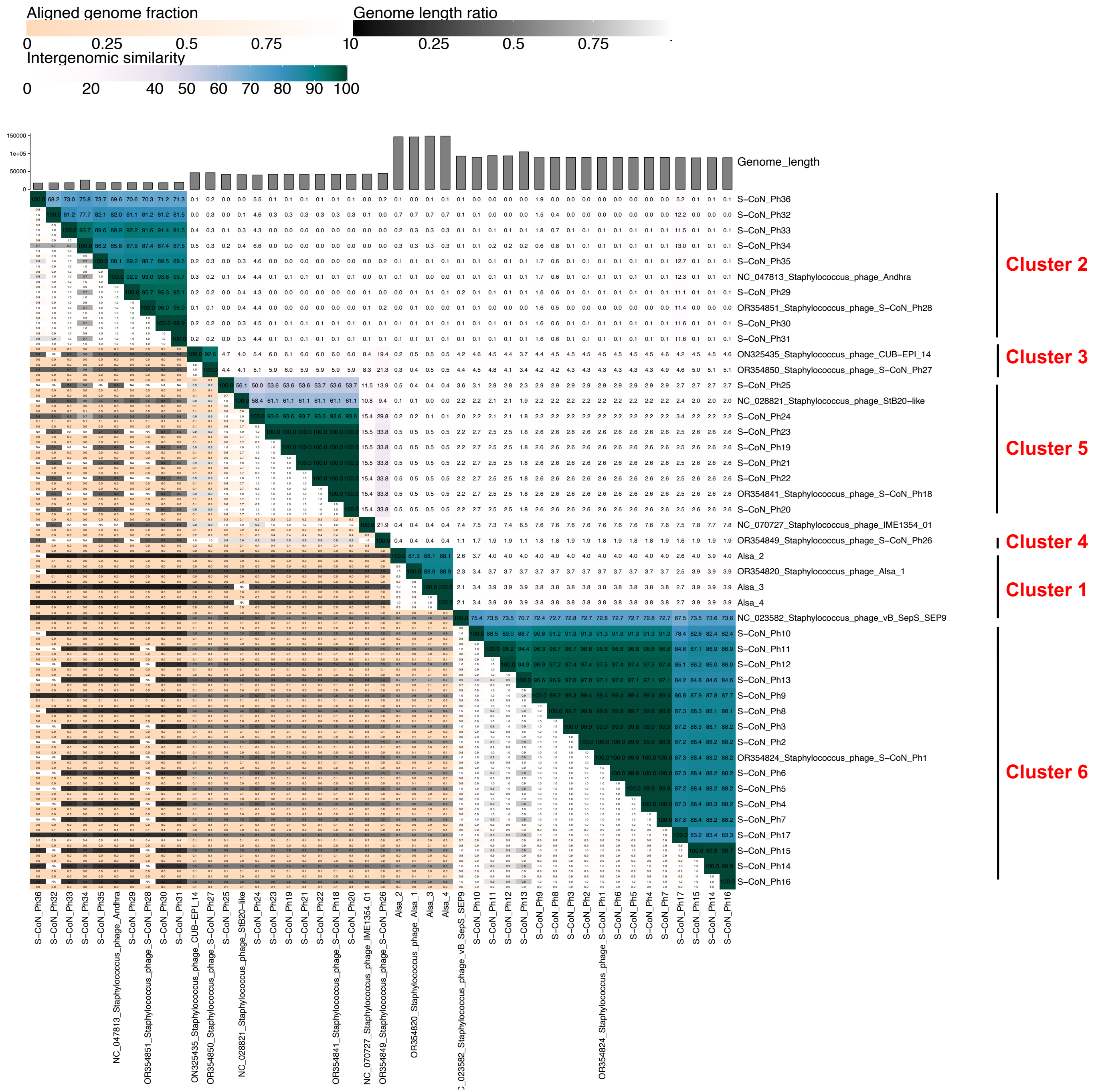


**Figure S1: Intergenomic similarities analysis of CoNS phages**

Heatmap generated using VIRIDIC to compare and cluster the 40 CoNS phages of this study. Intergenomic similarity between each pair of phage genomes is indicated on the right of the heatmap with values giving the similarity value (%) of each genome pair and blue-green shade gradient representing the degree of similarity. For each phage genome pair, three values are shown from top to bottom: aligned fraction of the first genome (in a row), the genome length ratio, and the aligned fraction of genome (column), respectively. Darker colour shows lower values indicating smaller aligned genome pairs (Orange to white colour range) or a large difference in the length of the genome pair (black to white scale). Published phage genomes were included as a reference for clusters 2-6 (cluster 2: NC_047813 ⌀Andhra; cluster 3: ON325435 ⌀CUB-EPI_14; cluster 4: NC_070727 ⌀IME1354_01; cluster 5: NC_028821 ⌀StB2—like; cluster 6: NC_023582 ⌀vB_SepS-SEP9). Staphylococcus phage IME1354_01 is a distinct species from ⌀ S-CoN_Ph26 (cluster 4) based on 95% definition of species.

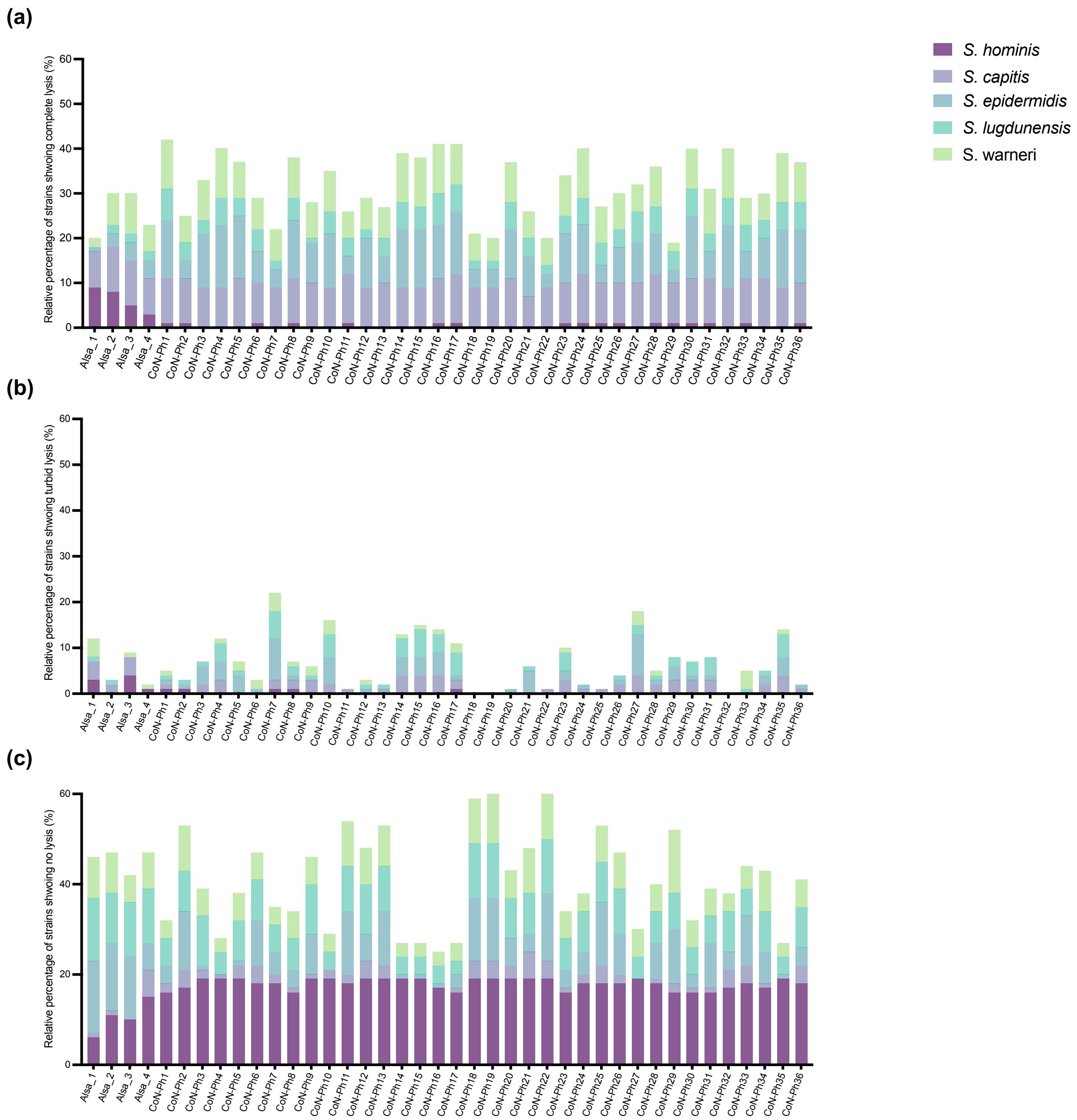


**Figure S2: Comparison of host species lysis.** From the host range assay spot assays, the relative percentage of each host strain with complete lysis zones after phage infection is shown for each of the 40 CoNS phages. CoNS hosts are indicated by the colours shown in the key.

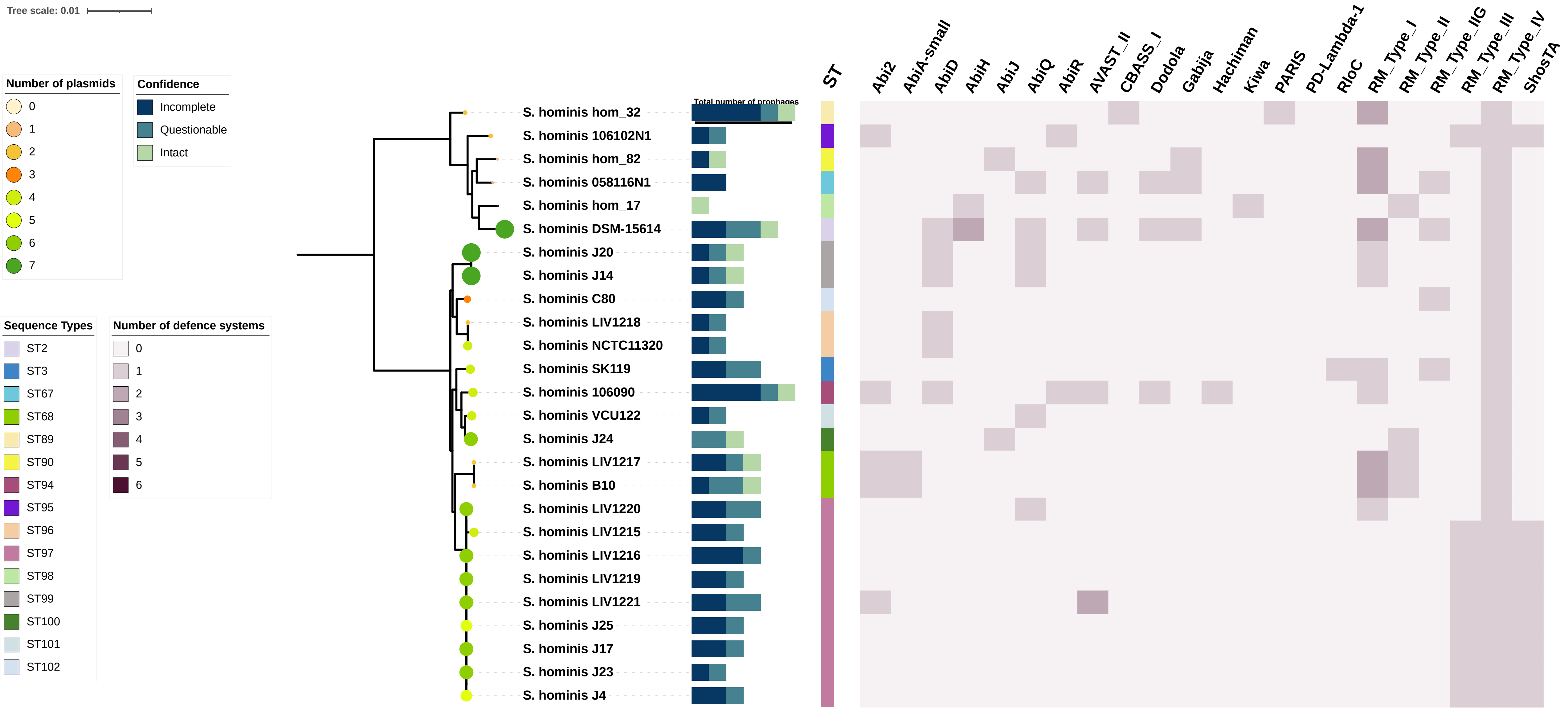


**Figure S3: Phage defence systems and plasmid content of *S. hominis.*** Phage defence systems in genomes of the *S. hominis* strains were detected using DefenseFinder. Number of defence systems present is scored (0-2 max from the key) and the total number of prophages (0-6) are indicated as either intact, incomplete, or questionable. Phylogenetic tree of *S. hominis* strains was generated using IQ-TREE. Sequence type (ST) of strains is indicated by vertical-coloured bars. Plasmid content (0-7 plasmids) is shown by different sized and coloured circle symbols, determined by PlasmidFinder.

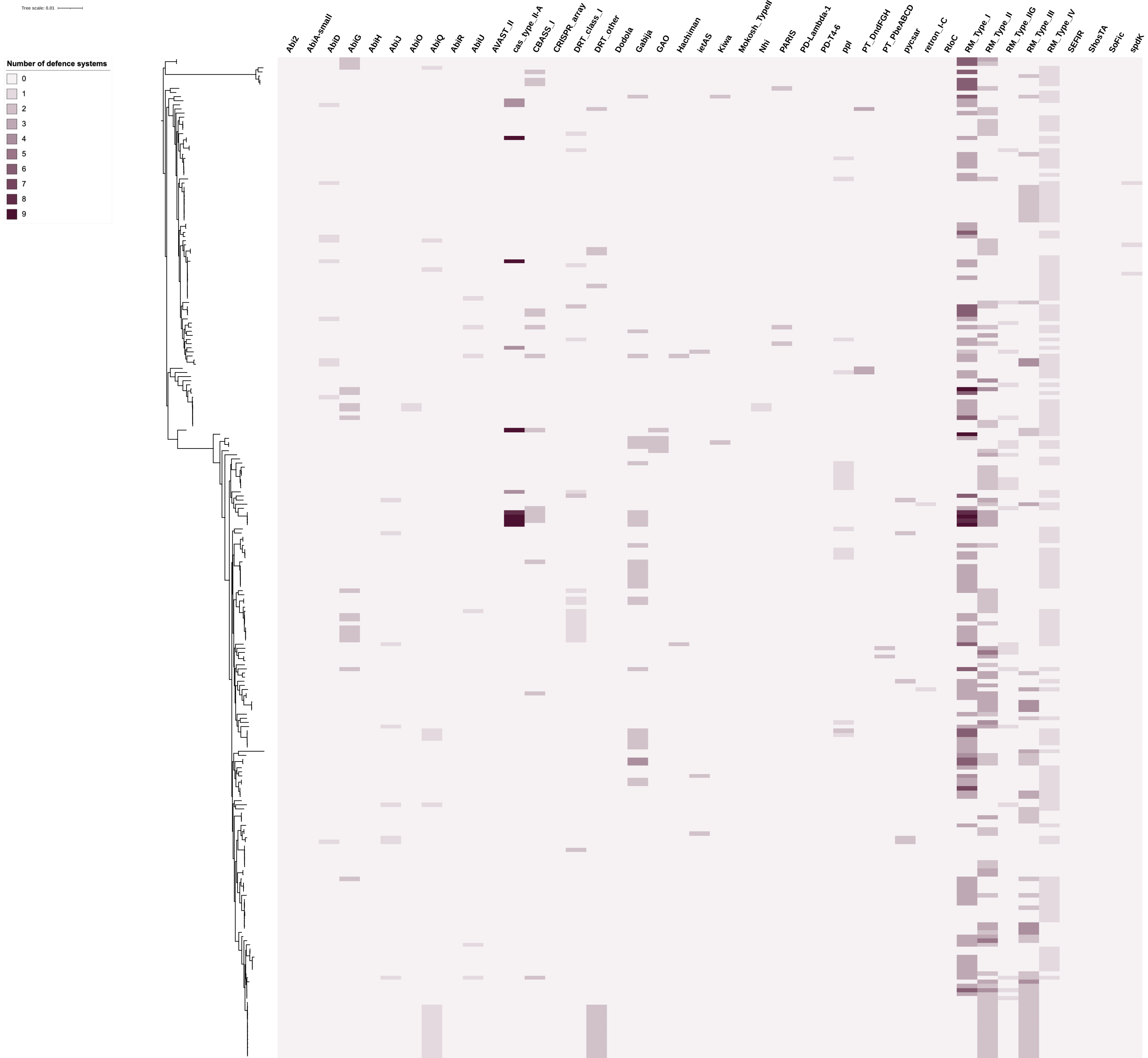


**Figure S4: Antiviral defence systems of publicly available *S. hominis* genomes.** Phage defence systems of 243 *S. hominis* strain genomes from NCBI database were determined using PADLOC. Number of defence systems present is scored (0-9).

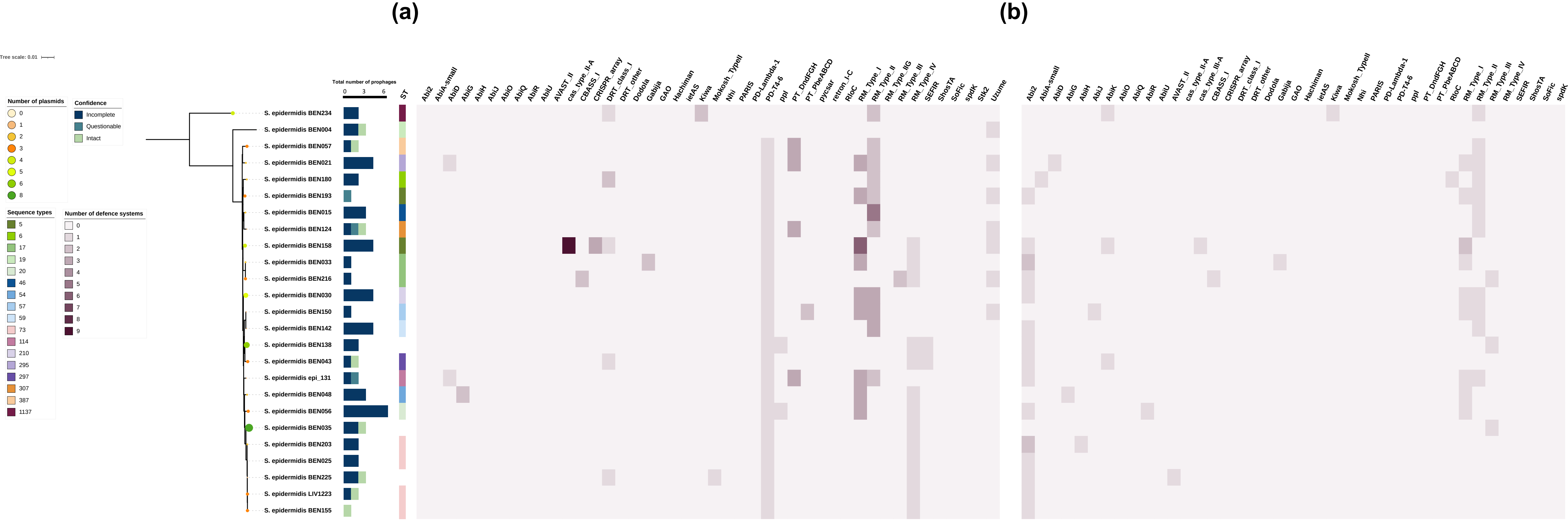


**Figure S5: Phage defence systems and plasmid content *S. epidermidis*.** Phage defence systems present in the genomes of *S. epidermidis* strains were detected using (a) PADLOC and (b) DefenseFinder. Number of defence systems present is scored (0-9), and the total number of prophages (0-6) are indicated as either intact, incomplete, or questionable. Phylogenetic tree of *S. epidermidis* strains was generated using IQ-TREE. Sequence type (ST) of strains is indicated by vertical-coloured bars. Plasmid content (0-8 plasmids) is shown by different sized and coloured circle symbols, determined by PlasmidFinder

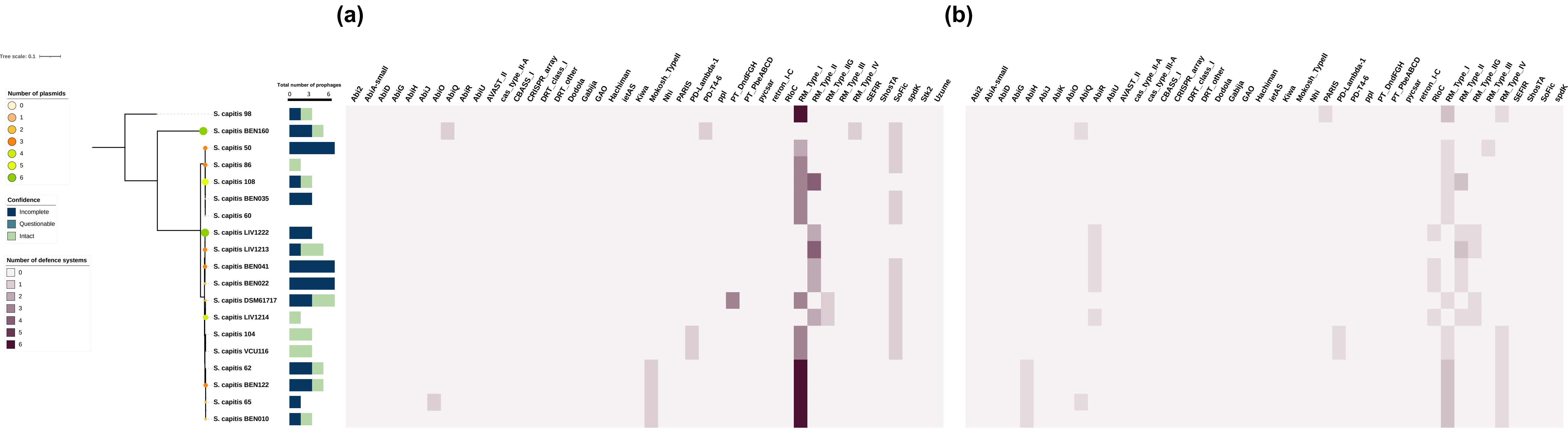


**Figure S6: Phage defence systems and plasmid content *S. capitis.*** Phage defence systems present in the genomes of *S. capitis* strains were detected using (a) PADLOC and (b) DefenseFinder. Number of defence systems present is scored (0-6), and the total number of prophages (0-6) are indicated as either intact, incomplete, or questionable. Phylogenetic tree of *S. capitis* strains was generated using IQ-TREE. Plasmid content (0-6 plasmids) is shown by different sized and coloured circle symbols, determined by PlasmidFinder
